# Neural markers of speech comprehension: measuring EEG tracking of linguistic speech representations, controlling the speech acoustics

**DOI:** 10.1101/2021.03.24.436758

**Authors:** Marlies Gillis, Jonas Vanthornhout, Jonathan Z. Simon, Tom Francart, Christian Brodbeck

**Affiliations:** ExpORL, Department of Neurosciences, KU Leuven, 3000 Leuven, Belgium; Department of Electrical & Computer Engineering; Department of Biology; Institute for Systems Research, University of Maryland, College Park, MD 20740, USA; Department of Psychological Sciences, University of Connecticut, Storrs, CT 06269

## Abstract

When listening to speech, our brain responses time-lock to acoustic events in the stimulus. Recent studies have also reported that cortical responses track linguistic representations of speech. However, tracking of these representations is often described without controlling for acoustic properties. Therefore, the response to these linguistic representations might reflect unaccounted acoustic processing rather than language processing. Here, we evaluated the potential of several recently proposed linguistic representations as neural markers of speech comprehension. To do so, we investigated EEG responses to audiobook speech of 29 participants (22 ♀). We examined whether these representations contribute unique information over and beyond acoustic neural tracking and each other. Indeed, not all of these linguistic representations were significantly tracked after controlling for acoustic properties. However, phoneme surprisal, cohort entropy, word surprisal, and word frequency were all significantly tracked over and beyond acoustic properties. We also tested the generality of the associated responses by training on one story and testing on another. In general, the linguistic representations are tracked similarly across different stories spoken by different readers. These results suggests that these representations characterize processing of the linguistic content of speech.

**Significance Statement:** For clinical applications it would be desirable to develop a neural marker of speech comprehension derived from neural responses to continuous speech. Such a measure would allow for behaviour-free evaluation of speech understanding; this would open doors towards better quantification of speech understanding in populations from whom obtaining behavioral measures may be difficult, such as young children or people with cognitive impairments, to allow better targeted interventions and better fitting of hearing devices.

## Introduction

When listening to natural running speech, brain responses time-lock to certain features of the presented speech. This phenomenon is called neural tracking (for a review, see, e.g., Brodbeck and Simon, 2020). Commonly, neural tracking is studied using an acoustic representation of the speech, for example, the envelope or spectrogram (Aiken and Picton, 2008; Ding and Simon, 2012b). Neural tracking of acoustic speech representations is modulated by attention (e.g., Ding and Simon, 2012a; Horton et al., 2014; O’Sullivan et al., 2015; Das et al., 2016) and speech understanding (Etard and Reichenbach, 2019; Iotzov and Parra, 2019; Vanthornhout et al., 2018; Lesenfants et al., 2019). However, the observation of neural speech tracking does not guarantee speech intelligibility, since music (Tierney and Kraus, 2014), and the ignored talker in the two-talker scenario, are also significantly tracked by the brain (Horton et al., 2014; O’Sullivan et al., 2015; Ding and Simon, 2012a).

A more promising avenue of neurally predicting behavioral speech understanding comes from recent studies which reported that linguistic properties, derived from the linguistic content of speech, are also tracked by the brain (Broderick et al., 2018; Brodbeck et al., 2018; Weissbart et al., 2020; Koskinen et al., 2020). Neural tracking of linguistic representations has mainly been studied with measures that quantify the amount of new linguistic information in a word, such as word surprisal or semantic dissimilarity. These representations show a negativity with a latency of around 400 ms relative to word onset (Broderick et al., 2018; Weissbart et al., 2020; Koskinen et al., 2020) which is in broad agreement with results of studies investigating the N400 event-related brain potential (ERP) response, an evoked brain responses to words, typically studied in carefully controlled stand-alone sentence or word paradigms (Frank et al., 2015; Frank and Willems, 2017; for a review on the N400 response, see e.g., Kutas and Federmeier, 2011 and Lau et al., 2008). Neural tracking of linguistic properties is also seen at the level of phonemes (Brodbeck et al., 2018; Gwilliams and Davis, 2020; Donhauser and Baillet, 2020). Importantly, several studies investigating neural tracking of linguistic representations report an absence of corresponding responses to the ignored speaker in a two-talker speech mixture, suggesting that these linguistic representations might selectively reflect speech comprehension (Brodbeck et al., 2018; Broderick et al., 2018).

Few studies, however, analyze neural tracking of linguistic representations while controlling for neural tracking of the acoustic properties of the speech (though see Brodbeck et al., 2018; Koskinen et al., 2020). This is problematic as linguistic features are often correlated with acoustic features. Indeed, Daube et al. (2019) found that acoustic features of speech can explain apparent responses to linguistic phoneme categories. Thus, without controlling for acoustic properties, speech tracking analysis might be biased to find spurious significant linguistic representations.

In addition to acoustic and linguistic representations, it is important to account for responses related to speech *segmentation*. These represent words or phonemes as discrete events, distinct from acoustic onsets, although the two are likely often correlated. Word onsets in continuous speech are associated with a characteristic brain response that is not purely acoustic (Brodbeck et al., 2018; Sanders and Neville, 2003). Therefore, in this study, we control for both acoustic features and speech segmentation, to identify the *added value* of linguistic representations.

Previous studies discuss one or a small number of linguistic representations separately, often without controlling for acoustic properties of the speech. Here, we combine recently proposed linguistic representations. We categorize these linguistic representations into three types depending on how they can contribute to language understanding: (a) phoneme level: phoneme surprisal and cohort entropy (Brodbeck et al., 2018), (b) word level: word surprisal, word entropy, word precision and word frequency (Weissbart et al., 2020; Koskinen et al., 2020) and (c) contextual level: semantic dissimilarity (Broderick et al., 2018).

In this study, we aim to assess the feasibility of linguistic representations as neural markers of speech comprehension in three ways. (1) We verify whether existing linguistic representations are tracked after controlling for the neural tracking of acoustic and speech segmentation properties. (2) Moreover, each linguistic representation should contribute unique information over and beyond other linguistic representations. (3) Finally, we examine whether the processing of these linguistic representations is generalizable across different stories. If so, these linguistic representations likely reflect linguistic speech processing and, therefore, would be good candidates for a neural marker of speech comprehension.

## Materials and Methods

### Participant Details

The electroencephalography (EEG) data of 29 young normal-hearing individuals (22 ♀) were analysed. The data were originally collected for other studies (Accou et al., 2020; Monesi et al., 2020). Participant age varied between 18 and 25 years old (mean*±*std= 20.81 *±* 1.94 years). The inclusion criteria were being a native speaker of Dutch and having normal hearing, which was verified using pure tone audiometry (octave frequencies between 125 and 8000 Hz; no hearing threshold exceeded 20 dB hearing level). The medical ethics committee of the University Hospital of Leuven approved the experiments, and all participants signed an informed consent form before participating (S57102).

### Experimental Procedure

#### EEG Experiment

##### Data acquisition

The EEG recording was performed in a soundproof booth with Faraday cage (at ExpORL, Dept. Neurosciences, KU Leuven) using a 64-channel BioSemi ActiveTwo system (Amsterdam, Netherlands) at a sampling frequency of 8192 Hz.

##### Stimuli presentation

Each participant listened to five Dutch stories: De kleine zeemeermin (DKZ), De wilde zwanen (DWZ), De oude lantaarn (DOL), Anna en de vorst (AEDV) and Eline (Table 1) presented in random order. Stories longer than 20 minutes were divided into parts, each lasting 13 to 15 minutes (DWZ and AEDV were divided into 2 parts, DKZ into 3 parts). One or two randomly selected stories or story parts were presented in noise, but for this study only participants who listened to all 3 parts of DKZ without background noise were included. Additionally, when testing the DKZ-based model on any of the other stories, only participants who listened to that story without noise were included (the resulting number of participants is summarized in Table 2).

**Table 1:** Details on the presented stories.

| Story | Author | Speaker | Duration (min) |
| --- | --- | --- | --- |
| De kleine zeemeermin (DKZ) | H. C. Andersen | Katrien Devos (♀) | 46.08 |
| De wilde zwanen (DWZ) | H. C. Andersen | Katrien Devos (♀) | 27.46 |
| De oude lantaarn (DOL) | H. C. Andersen | Katrien Devos (♀) | 16.02 |
| Anna en de vorst (AEDV) | Unknown | Wivine Decoster (♀) | 25.51 |
| Eline | Rascal | Luc Nuyens (♂) | 13.33 |

**Table 2:** Amount of participants used for the across story comparisons.

| DWZ part 1 | DWZ part 2 | DOL | AEDV part 1 | AEDV part 2 | Eline |
| --- | --- | --- | --- | --- | --- |
| 20/29 | 19/29 | 22/29 | 15/29 | 23/29 | 23/29 |

Participants were instructed to attentively listen to the presented story. They were notified beforehand that content-related questions are asked at the end of the story to motivate them to listen to the story actively.

The speech stimuli were presented bilaterally at 65 dB sound pressure level (SPL, A-weighted) through ER-3A insert earphones (Etymotic Research Inc, IL, USA) using the software platform APEX (Dept. Neurosciences, KU Leuven) (Francart et al., 2008).

### Signal Processing

#### Processing of the EEG signals

The EEG recording with a sampling frequency of 8192 Hz was downsampled to 256 Hz to decrease the processing time. We filtered the EEG using a multi-channel Wiener filter (Somers et al., 2018) to remove artifacts due to eye blinks. We referenced the EEG to the common-average and filtered the data between 0.5 and 25 Hz using a Chebyshev filter (Type II with an attenuation of 80 dB at 10% outside the passband). Then additional downsampling to 128 Hz was done.

#### Extraction of the predictor variables

In this study, we combined acoustic speech representations with recent proposed linguistic representations. We used speech representations for acoustic properties of the speech (spectrogram, acoustic onsets), segmentation of the speech (phoneme onsets, word onsets, function word onsets and content word onsets) and linguistic properties (phoneme surprisal, cohort entropy, word surprisal, word entropy, word precision, word frequency, semantic dissimilarity). An example of these speech representations is visualized in Figure 1 (for illustration purposes, only one band of the 8-band spectrogram and acoustic onsets is visualized).

**Figure 1:**
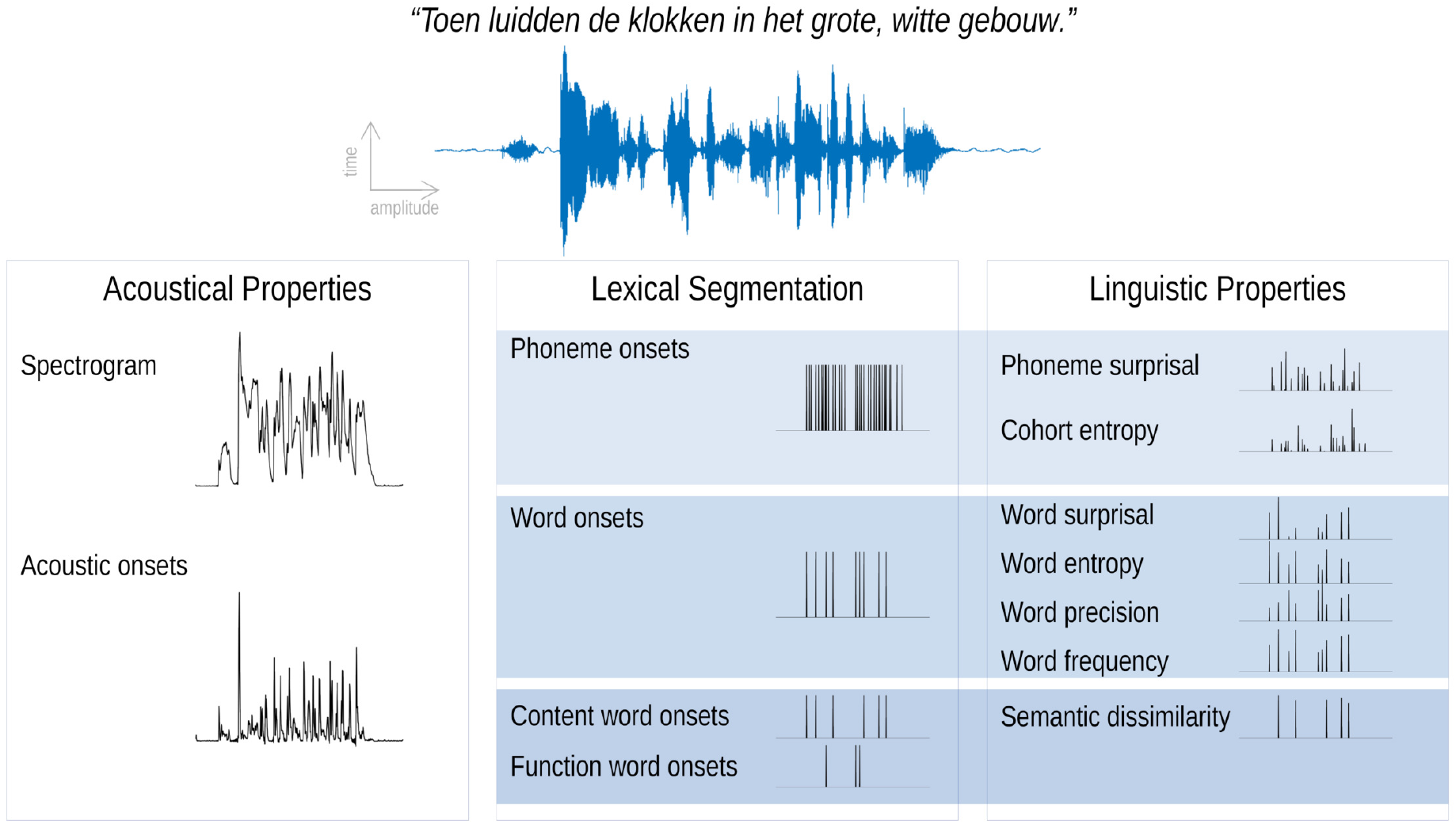
Speech representations used in this study. For illustration purpose, only one band of the spectrogram and acoustic onsets is visualized.

##### Spectrogram and acoustic onsets

Both of these speech representations reflect the continuous acoustic power of the presented speech stimuli. A spectrogram representation was obtained using the Gammatone Filterbank Toolkit 1.0 (Heeris (2014); frequency cut-offs at 20 and 5000 Hz, 256 filter channels and a window time of 0.01 second). This toolkit calculates a spectrogram representation based on a series of gammatone filters inspired by the human auditory system (Slaney, 1998). The resulting filter outputs with logarithmic center frequencies were averaged into 8 frequency bands (frequencies below 100 Hz were omitted similar to Brodbeck et al. (2020)). Additionally, each frequency band was scaled with exponent 0.6 (Biesmans et al., 2016) and downsampled to the same sampling frequency as the processed EEG, namely 128 Hz.

For each frequency band of the spectrogram, an acoustic onsets representation was computed by applying an auditory edge detection model (Fishbach et al., 2001) (using a delay layer with 10 delays from 3 to 5 ms, a saturation scaling factor of 30 and receptive field based on the derivative of a Gaussian window with a standard deviation of 2 ms (Brodbeck et al., 2020)).

##### Phoneme onsets and word onsets

Time-aligned sequences of phonemes and words were extracted by performing a forced alignment of the identified phonemes using the speech alignment component of the reading tutor (Duchateau et al., 2009). The resulting representations were one-dimensional arrays with impulses on the onsets of, respectively, phonemes and words.

##### Content word onsets and function word onsets

The Stanford Parser (Klein and Manning, 2003b,a) was used to identify the part-of-speech category of each word. We subsequently classified the words into 2 classes: (a) open class words, also referred to as content words, which included all adjectives, adverbs, interjections, nouns and verbs and (b) closed class words, also referred to as function words, which included all adpositions, auxiliary verbs, conjunctions, determiners, numerals, articles and pronouns. The resulting representations were one-dimensional arrays with impulses at the onsets of, respectively, content or function words.

##### Linguistic representations at the phoneme level

Two linguistic phoneme representations were modeled to describe each phoneme’s informativeness in its lexical context, namely *phoneme surprisal* and *cohort entropy* (Brodbeck et al., 2018). Both representations are derived from the active cohort of words (Marslen-Wilson, 1987): a set of words which start with the same acoustic input at a given point during the word. Phoneme surprisal reflects how surprising a given phoneme is, given the previous phonemes. It is calculated as the negative logarithm of the inverse conditional probability of each phoneme given the preceding phonemes in the word. Cohort entropy reflects the degree of competition among words which are compatible with the partial phoneme string from word onset to the current phoneme. It is expressed as the Shannon entropy of the active cohort of words at each phoneme (for details of both representations, see Brodbeck et al. (2018)). The lexicon for determining the cohort was based on a custom pronunciation dictionary maintained at our lab (created manually and using grapheme-to-phoneme conversion; containing 9157 words). The prior probability for each word was based on its frequency in the SUBTLEX-NL database (Keuleers et al., 2010). Phoneme surprisal and cohort entropy were calculated from this cohort model according to the equations below. The initial phoneme of each word was not modeled in these representations. The resulting representations were one-dimensional arrays with impulses at phoneme onsets modulated by the value of respectively surprisal or entropy, except for the word’s initial phoneme.

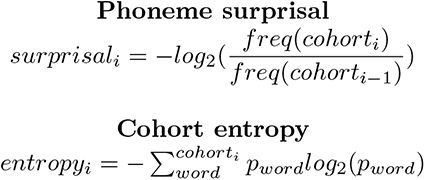

##### Linguistic representations at the word level

Linguistic word representations were derived using a Dutch 5-gram model (by Verwimp et al. (2019) using the corpora: Corpus of Spoken Dutch (Oostdijk et al., 2000) and a database of subtitles). N-gram models are Markov models which describe a word’s probability based on its *n −* 1 previous words. This way, it allows describing each word’s informativeness independent of sentence boundaries. Here, we focused on *word surprisal*, *word entropy*, *word precision* and *word frequency*.

Word surprisal was calculated as the negative logarithm of the conditional probability of the considered word given the four preceding words. It represents how surprising a word is given the four preceding words.

Word entropy is the Shannon entropy of the word given the four preceding words. It reflects the degree of competition between the word possibilities. A higher word entropy reflects that more words have a high probability of occurring after the four previous words.

Word precision was defined as the inverse of the word entropy. A high word precision indicates that only a few words are candidates to follow the four previous words. Therefore, the word can be predicted with high precision.

Word frequency was included as the negative logarithm of the word’s unigram probability. It represents word probability independent of the preceding words. Please note the negative logarithm: words with a high frequency yield a low value and vice versa. Note that some of the methods differ slightly between phoneme-and word-level representations; we opted to use representations as close as possible to those used previously in the literature.

The resulting representations were one-dimensional arrays with impulses at word onsets modulated by the value of, respectively, surprisal, entropy, precision or word frequency.

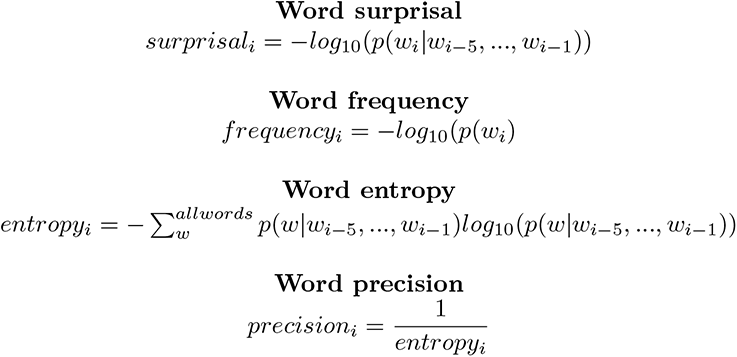

##### Semantic representation

To describe the influence of semantic context, *semantic dissimilarity* was used as a measure of how dissimilar a content word is compared to its preceding context (Broderick et al., 2018). Unlike linguistic representations at the word level, this representation does take into account sentence boundaries. For each content word in the story, a word embedding was retrieved from a database with word embeddings obtained with word2vec (Tulkens et al., 2016) using a combination of different Dutch text corpora (Roularta (Roularta Consortium, 2011), Wikipedia (Wikipedia, 2015), SoNaR corpus (Oostdijk et al., 2013)). To obtain a value of semantic dissimilarity for a content word, the word embedding of the considered word was correlated (Pearson’s correlation) with the average of the previous content words in the considered sentence. This correlation value was subtracted from 1 to obtain a value which reflects how dissimilar the word is compared to its context. If the word was the initial content word of the sentence, its word embedding was correlated with the average of the word embeddings of the content words in the previous sentence. The resulting representation was a one-dimensional array with impulses at content word onsets modulated by the value of how dissimilar the considered content word is compared to its context.

#### Determination of neural tracking

We focused on a linear forward modelling approach that predicts the EEG response given some preceding speech representations. This forward modelling approach results in (a) a temporal response function (TRF) and (b) a prediction accuracy for each EEG channel. A TRF is a linear kernel which describes how the brain responds to the speech representations. This TRF can be used to predict the EEG-response by convolving it with the speech representations. The predicted EEG-response is then correlated with the actual EEG-response, and correlation values are averaged across EEG channels to obtain a single measure of prediction accuracy.

This prediction accuracy is seen as a measure of neural tracking: the higher the prediction accuracy, the better the brain tracks the stimulus.

(a) To estimate the TRF, we used the Eelbrain toolbox (Brodbeck, 2020) which estimates a TRF for each EEG electrode separately using the boosting algorithm (David et al., 2007). We used 4-fold cross-validation (4 equally long folds; 2 folds used for training, 1 for validation and 1 fold unseen during training for testing; for each testing fold, 3 TRF models were fit, using each of the remaining 3 folds as the validation fold in turn). Cross-validation employing the additional test stage using unseen data allows a fair comparison between models with different numbers of speech representations. TRFs covered an integration window from 0 to 900 ms (with a basis of 50 ms Hamming windows, and selective stopping based on the *R*_2_-norm after 1 step with error increase). For analyzing the TRFs, the resulting TRFs were averaged across all folds. (b) To calculate the prediction accuracy, the average TRF from 3 complimentary training folds was used to predict the corresponding unseen testing fold. Predictions for all testing folds were then concatenated to compute a single model fit metric. The correlation between the predicted and actual EEG was averaged across channels to obtain the prediction accuracy.

To evaluate whether a speech representation had a significant added value, we compared whether the prediction accuracy significantly increased when the representation was added to the model (e.g., to determine the added value of word onsets over the spectrogram, we compared the prediction accuracy obtained with the model based on the spectrogram to the prediction accuracy of the model based on a combination of the spectrogram and word onsets).

In sum, we investigated which linguistic speech representations are significantly tracked by the brain by examining whether the prediction accuracy significantly improves when representations are added. If so, we investigated the neural response to the linguistic speech representations by examining the TRFs.

#### Determination of the peak latency

The latencies of the response peaks in TRFs were determined for the linguistic speech representations at the phoneme level (Brodbeck et al., 2018). Based on the mean TRFs across participants, we identified different time windows in which we determined the peak latency (30-90 ms, 90-180 ms and 180-300 ms). For each subject, the latency was determined as the time of the maximum of the absolute values of the TRF across channels.

#### Statistical analysis

We calculated the significance level of these prediction accuracies by correlating the EEG responses with EEG-shaped noise (e.g., noise with the same frequency spectrum as the EEG responses), which was done 1000 times for each participant. The significance level was determined as the 97.5th percentile of the obtained correlations. This resulted in a significance level for each subject. The maximal significance level was 0.0019 for a correlation average across all channels, and 0.003 for an individual channel. All obtained prediction accuracies averaged across channels exceeded this significance level. Therefore, the significance of the prediction accuracies is not explicitly mentioned in the remainder of this manuscript.

For univariate statistical analysis, we used the R software package (version 3.6.3) (R Core Team, 2020). We performed one-sided Wilcoxon signed-rank tests to identify whether the linguistic representations had added value beyond acoustic and speech segmentation representations. The outcomes of such a test are reported with a p-value and effect size. All tests were performed with a significance level of *α* = 0.05. Effect sizes are derived from the z-scores underlying the p-values, divided by the root of the number of observations as proposed by Field et al. (2012). This method allows approximating the effect size measure for non-parametric tests, which is similar to Cohen’s d. Effect sizes above 0.5 indicate a large effect. To inspect whether the latencies differed significantly, a two-sided Wilcoxon signed-rank test was performed.

To compare topographic responses, we applied a method proposed by McCarthy and Wood (1985), which evaluates whether the topography differs between two conditions when amplitude effects are discarded. The method is based on an ANOVA testing for an interaction between sensor and condition, i.e., testing whether the normalized response pattern across sensors is modulated by condition. We compared the average topographic response within specific time windows. These time windows were determined as the intersections of the time intervals in which the smoothed average TRFs of a frontal and a central channel selection significantly differed from 0, for a duration of more than one sample, after smoothing using a hamming kernel of 100 ms. This smoothing was performed to decrease the inter-subject variability of the peak latencies. A significant difference in the topography suggests that a different neural source configuration evokes the two topographies. A different neural source configuration implies that either different neural sources are active or the relative strength of these neural sources has changed.

To determine the significance of TRFs, we used mass-univariate cluster-based permutation tests as recommended by (Maris and Oostenveld, 2007), using the Eelbrain (Brodbeck, 2020) implementation. This method first calculates a univariate t-statistic at each time point and sensor, and then finds spatio-temporal clusters in which this statistic exceeds a certain value (we used a cluster-forming threshold of uncorrected p=0.05). For each cluster, the cluster-mass statistic is computed, which is equal to the sum of all t-values in the cluster. To calculate a p-value, this cluster-mass is then compared to a null distribution based on the largest cluster-mass in 10,000 permutations of the data (for one-sample tests: random sign flips; for related measures t-tests: random permutation of condition labels). We tested whether the average TRF was significantly different from 0 using permutation tests based on two-tailed one-sample t-tests (by permuting the sign of the values). To determine whether the TRF differed between two speech representations, we used permutation tests based on related-measures t-tests (by permuting the values between the different speech representations). For determining significant clusters, we used a corrected significance level of *α* = 0.05.

We also compared how well responses of the different stories could be predicted using the same TRFs. Among the different stories, we noticed that the added value of the linguistic representations varied. Therefore, we investigated which predictors could explain this variance among the different stories. To do so, we used the Buildmer toolbox to identify the best linear mixed model (LMM) given a series of predictors based on the likelihood-ratio test (Voeten, 2020). The analysis included a factor with a level for each story, a continuous predictor reflecting the presentation order, a distance-from-training-data metric and a random effect for participant. The presentation order predictor reflects the linear presentation order during the experiment and would therefore be able to model changes of neural tracking over the course of the experiment. The distance-from-training-data metric is calculated as the number of stories presented between the presentation of the story and DKZ. Hereafter referred to as presentation distance. This metric would allow to investigate whether neural tracking is affected by the subject’s mental state (e.g. tiredness; stories presented right before or after the training story DKZ have a similar mental state and therefore the neural tracking should be similar).

Moreover, we reported specific speech characteristics of each story: duration (in minutes), word count, word rate (defined as word count/duration), voice frequency (defined as the frequency with maximal power), speaker’s sex (Table 3).

**Table 3:** Predictors based upon the speech characteristics for each story.

| story | speaker's sex | word count | duration | word rate | voice frequency | p-value of wilcoxon test |
| --- | --- | --- | --- | --- | --- | --- |
| DKZ_1 | F | 2257 | 15.1 | 149.47 | 142 | NA |
| DKZ_2 | F | 2192 | 15.15 | 144.69 | 141 | NA |
| DKZ_3 | F | 2279 | 15.87 | 143.63 | 140 | NA |
| DOL | F | 2365 | 16.03 | 147.51 | 139 | n.s. |
| DWZ_1 | F | 1969 | 13.88 | 141.82 | 240 | n.s. |
| DWZ_2 | F | 1879 | 13.88 | 135.34 | 238 | p=0.03 |
| AEDV_1 | F | 1708 | 13.17 | 129.72 | 183 | p<0.001 |
| AEDV_2 | F | 1669 | 12.68 | 131.59 | 178 | p=0.013 |
| Eline | M | 1935 | 13.55 | 142.81 | 116 | p<0.001 |

## Results

### Linguistic properties are reliably tracked within story

The analysis of this section followed three steps, with each subsequent step based on the results of the previous step.

1. We combined the linguistic representations at each level to identify whether the level as a whole contained representations which significantly improved the predictions. For each level, we determined whether the combination of the different linguistic representations significantly increased the prediction accuracies if added to a model with acoustic and speech segmentation properties of the speech.
2. We identified, at each level, which speech representations contributed unique information to the model. By comparing the prediction accuracies of a model with all linguistic representations of the considered level to the prediction accuracies of a model with a specific linguistic representation left out, we determined the added value of the left-out representation. If the prediction accuracy is significantly higher for the model which includes all linguistic representations, then the left-out linguistic representation contributes unique information to the model, over and beyond the other representations.
3. We verified whether the significant representations at the different levels had an added value over and beyond each other, following a similar strategy as above.

We first analyzed responses to linguistic representations in a single story (DKZ: 46 minutes; 29 participants). At each level of representations, we first verified whether the full set at each level of linguistic representations had an added value over and beyond the acoustic and speech segmentation representations (added value visualized in Figure 2.A). At both the phoneme and the word level, a model which included all linguistic representations of the considered level showed a significantly higher prediction accuracy compared to a model which only included acoustic and speech segmentation representations (phoneme level: p<0.001, effect size=0.682; word level: p=0.015, effect size=0.405). However, semantic dissimilarity did not have a significant added value over and beyond acoustic and speech segmentation representations (p=0.641; Figure 2.A). In previous literature, significant neural tracking of semantic dissimilarity is reported, however, without controlling for acoustic feature or content word onsets. Consistent with this earlier result, we did observe that semantic dissimilarity by itself does yield prediction accuracies significantly above 0 (p<0.001; effect size=0.925). We further found that semantic dissimilarity retains its added value over and beyond content word onsets (p<0.001, effect size=0.592). However, as stated above, when fully controlling for acoustic speech representations over and above word and phoneme onsets, no added value of semantic dissimilarity was observed.

**Figure 2:**
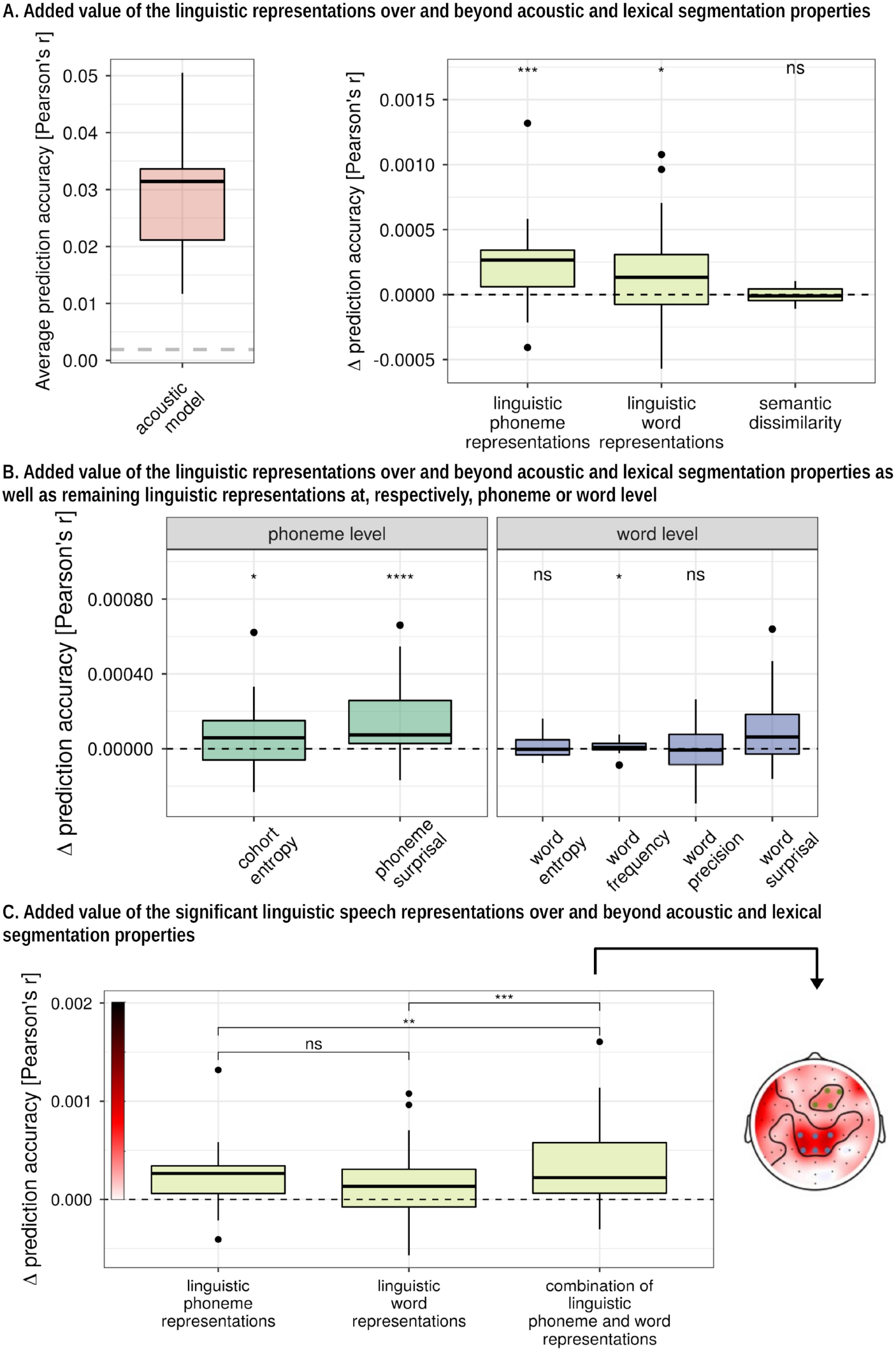
Added value of linguistic representations averaged across channels: Panel A: Raw prediction accuracies obtained with a model which includes only acoustic and speech segmentation properties (e.i. baseline model). The horizontal dashed grey line indicates the significance level of the prediction accuracies averaged across channels (left).Increase in prediction accuracy (Pearson’s r) of the combined representations at each level compared to a baseline model which included acoustic and speech segmentation properties of the speech (right). Panel B: Increase in prediction accuracy (Pearson’s r) of each representation compared to a baseline model which includes the other linguistic representations at the considered level. Panel C: Increase in prediction accuracy compared to a baseline model of a combination of the significant features averaged across channels (left) and in sensor space (right). *(*: p<0.05, **:p<0.01, ***: p<0.001, ****:p<0.0001)*

At the phoneme and word level, we determined which linguistic representations within the considered level contributed significantly over and beyond the other linguistic representations at that level, in addition to the acoustical and speech segmentation representations (Figure 2.B). At the phoneme level, phoneme surprisal and cohort entropy both had a significant added value over and beyond each other and acoustic speech representations (phoneme surprisal: p<0.001, effect size=0.702; cohort entropy: p=0.046, effect size=0.313). However, for the linguistic representations at the word level, only word surprisal (p=0.004, effect size=0.492) and word frequency (p=0.019, effect size=0.384) contributed significantly to the model while word entropy (p=0.275) and word precision (p=0.609) did not have an added value.

Subsequently, we combined all the significant linguistic representation at the word and phoneme levels derived from the first analysis. The significant linguistic speech representations at the phoneme level had an added value over and beyond the significant linguistic speech representations at the word level (p=0.001, effect size=0.589) and vice versa (p=0.008, effect size=0.448). On average, the prediction accuracy improved by

1.05 % when the linguistic representations were added to a model which only contains the acoustic and speech segmentation properties of the speech (prediction accuracy increased with 3.4 *×* 10*^−^*^4^, p<0.001, effect size=0.713; Figure 2.C). The increase in prediction accuracy over the different sensors is visualized in Figure 2.C (right inset).

### Neural responses to linguistic features

To investigate the neural responses to the linguistic features, we examined the TRFs within a channel selection where the linguistic representations significantly improved the model. First, we averaged across either a central or frontal channel selection, depicted in Figure 2.C (right). Second, we determined with cluster-based permutation tests whether the TRFs differed significantly from 0. This analysis indicates in which latency range, the considered linguistic representation shows a consistent neural response across subjects. Subsequently, within the intersection of these latency ranges (annotated with the grey horizontal bar in Figure 3) between two linguistic representations of the considered level, we investigated whether the topography significantly differed between the two linguistic representations using the method proposed by McCarthy and Wood (1985). A significant difference indicates that the two topographies differ in the underlying neural source configurations, rather than just in amplitude.

**Figure 3:**
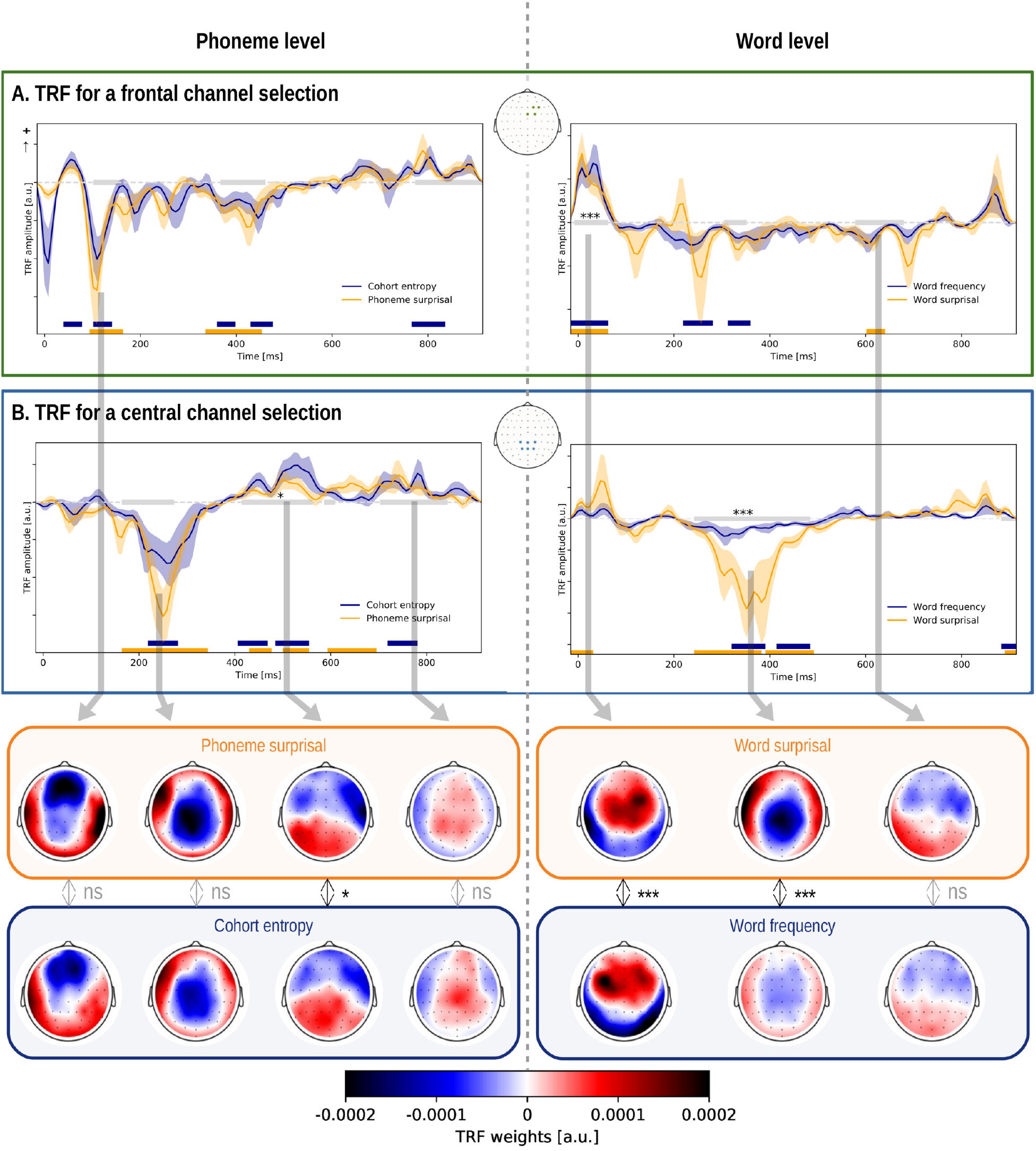
TRFs of linguistic representations at the phoneme and word level: The TRFs, averaged across participants for the different linguistic representations and a channel selection (shown in the central inset). The shaded area denotes the within-subject standard error of the average TRF. The time windows which are significantly different from zero are annotated with a horizontal line in the same colour as the TRF of the speech representations. The grey horizontal line denotes the time windows in which the two representations’ average topographies were compared. If a significant difference in topography was observed, the time window is annotated with a star. The corresponding topographies, averaged across this time window, are given as inset below encircled in the same colour as the TRF. The reported p-value is the result of the McCarthy-Wood method (for this method the normalized topographies are used which are not visualized here). *(*: p<0.05, **:p<0.01, ***: p<0.001, ****:p<0.0001)*

The TRFs for phoneme surprisal and cohort entropy are shown in Figure 3 (left) for both channel selections (TRFs for all channels are shown in Figure 3-2). Both linguistic representations at the phoneme level show a significant frontal negativity around 100 ms, and a significant central negativity around 250 ms followed by positive activity from 400 to 800 ms in central regions. We asked whether there is any evidence that the neural source configuration underlying the two TRFs is different. We did not observe a significant difference in topography of the earlier negativity around 100 ms (Figure 3: bottom; Table 3-2). Interestingly, we observed a significantly different topography in the time window from 414 ms to 562 ms, which indicates that the underlying neural source configuration is different (Figure 3; left; Figure 3-1; Table 3-1). However, judging from the difference map shown in Figure 3-1, the difference in topography is not easy to interpret, and could be due to a complex interplay between different neural sources. As the difference is difficult to interpret and the p-value is just below the significance threshold, the observed difference in topography might not be a robust effect.

The TRF to both phoneme-level representations shows 3 peaks (TRFs for all channels are shown in Figure 3-2a). Based upon the averaged TRF across participants, we identified 3 time windows wherein we determined the peak latency (respectively, 30 to 90 ms, 90 to 180 ms and 180 to 300 ms). We did not observe a significant difference in latency of all 3 peaks of phoneme surprisal and cohort entropy (30 to 90 ms: p=0.257, 90 to 180 ms: p=0.108, 180 to 300 ms: p=0.287).

The neural responses to the linguistic representations at the word level are shown in Figure 3 (right) (TRFs for all channels are shown in Figure 3-3). Both representations show a significant positive activation in frontal regions around 50 ms, and a prominent negativity around 300 to 400 ms after the word onsets. However, the amplitude of this negativity is smaller for word frequency. Interestingly, we identified a significant difference in topography for this negativity after discarding amplitudes effects (Figure 3: bottom; Figure 3-4; Table 3-3). The negativity for word frequency is situated more centrally compared to the negativity for word surprisal. The topography during the early responses to the word onset is also significantly different between the two speech representations (Figure 3-4; Table 3-4. Figure 3-4 shows that early activity of word surprisal shows more central activation while the early activity of word frequency is situated more laterally).

Additionally, we compared the topography of the negativity around 200 ms of phoneme surprisal (164 ms to 343 ms) to the topography of the negativity around 400 ms of word surprisal (242 ms to 531 ms). The method proposed by McCarthy and Wood (1985) did not identify a significant difference between these topographies (Figure 3-5). This suggests that the two effects reflect a shared neural process responding to surprising linguistic input.

### Neural processing of content and function words

Initially, we used a baseline model that represented acoustic properties and the speech segmentation. This model was kept constant to investigate the added value of different speech representations. However, as we did not observe an added value of semantic dissimilarity, which was encoded at every content word, we investigated whether word onsets split up depending on the word class had an added value (Brennan and Hale, 2019). In this analysis, we determined whether the differentiation between content and function words has an added value by three different models: (A) a baseline model including word onsets and the linguistic representations at the word level independent of the word class, (B) a model which differentiated between content words and function words for both word onsets as well as the linguistic representations at the word level, and (C) a model including a differentiation between content and function words for the word onsets but not for the linguistic representations at the word level. For the latter two models, a word onsets predictor for all words was included as well to capture TRF components shared between all words.

We observed an added value of the word class predictors (model C obtains higher prediction accuracies compared to model A: p < .001, effect size = 0.723; inset in Figure 4). However, we did not observe an added value of differentiating the linguistic speech representations at the word level depending on the word class (model B does not obtain higher prediction accuracies than model C: p=0.947). Thus, the response to function words differs from the response to content words, but the word class does not modulate responses related to word frequency and surprisal.

**Figure 4:**
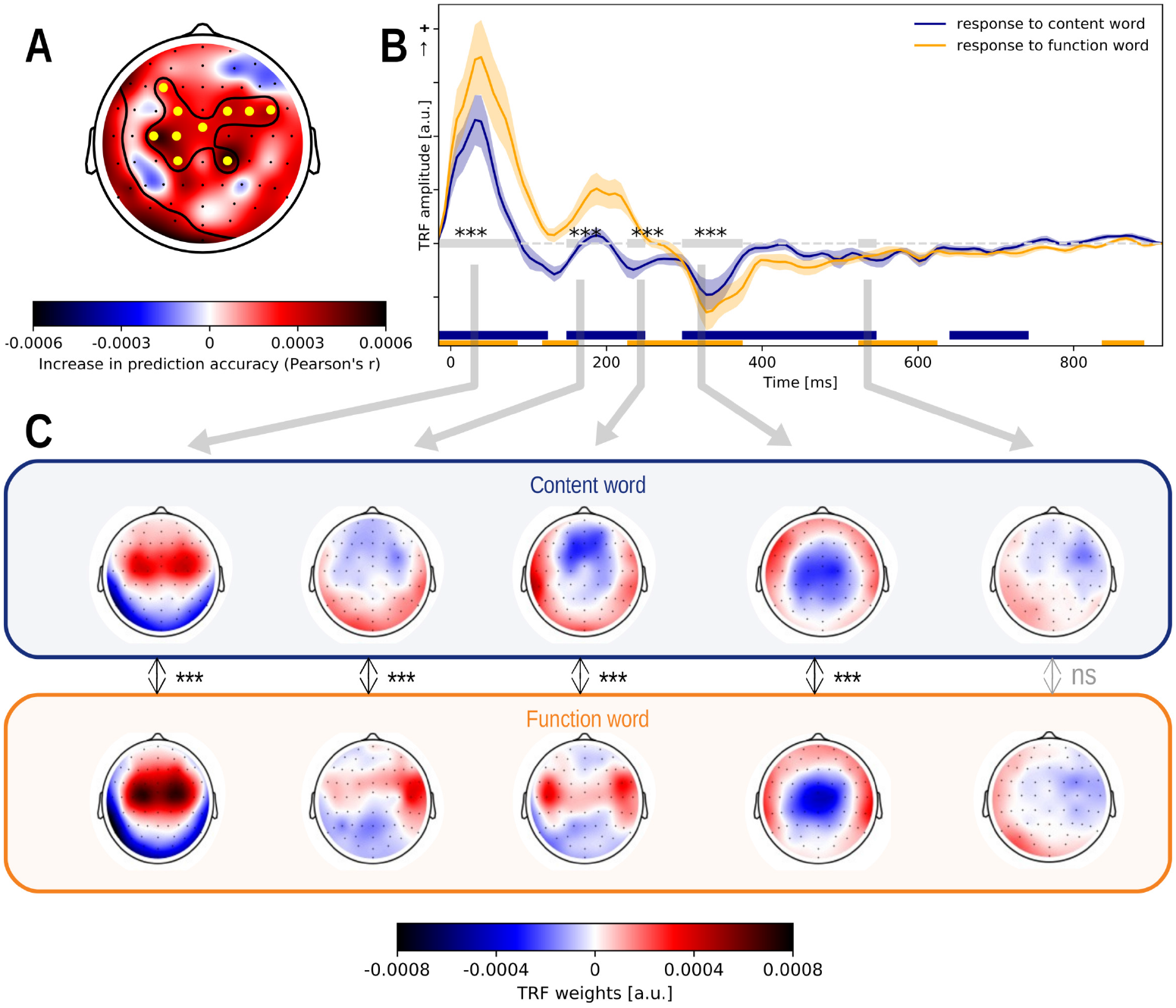
The response to words depends on the word class: Panel A: The increase in prediction accuracy (Pearson’s r) of including representations for the word classes into the model. Panel B: The response to content (blue) and function words (yellow), averaged across participants and a channel selection (marked yellow in Panel A) where the improvement of the differentiation between the word classes was significant. The shaded area denotes the standard error of the average TRF. The windows which are significantly different from zero are annotated with a horizontal line in the same colour as the TRF. The grey horizontal line denotes the time windows in which the two speech representations’ average topographies are compared. If a significant topography was observed, the time window is annotated with a grey star. The corresponding topographies averaged across this time window, are given as insets below encircled in the same colour as the TRF. The reported p-value is based on the McCarthy-Wood method (for this method the topographies were normalized which are not visualized here). *(*: p<0.05, **:p<0.01, ***: p<0.001, ****:p<0.0001)*

Subsequently, we investigated the difference in the response to content and function words by looking at the TRFs (Figure 4; TRFs for all channels are shown in Figure 4-2). For this analysis, we combined the TRF of word onsets and the TRF of content or function words to obtain the response to, respectively, a content or function word. The neural responses to words in both classes showed a significant central positivity around 50 ms and a negativity around 350 ms. In addition, the response to content words showed a significant positivity around 200 ms, while a slightly earlier significant negative response was observed in the response to function words.

For all the above mentioned time windows, a significant difference in topography was observed. The early response to function words is situated more centrally than the response to content words while the early response to content words shows more frontal activity. In the subsequent time windows around 200 ms, the response to content words shows a frontal negativity. The response to function words around 200 ms resembles the early response to word onsets with lateralized frontotemporal activation (see Figure 4.C, first topography). Around 350 ms, a central negativity is observed for both responses. This time window is also associated with a difference in topography, but the difference between the two topographies is difficult to interpret (difference is visualized in Figure 4-1; Table 4-1).

As noted above, the topography of the response to function words around 200 ms resembles the early ( 50 ms) response to all word onsets. This might be due to the properties of the different word classes: the duration of function words is generally shorter than that of content words which implies that the time interval between a word and its next word is shorter for function words (on average 239 ms for a function word while 600 ms for a content word). The response to function words might thus be more contaminated by a response to the subsequent word onset. To investigated whether the TRF of function words was contaminated by the onset of the next word, we divided function words into two categories: function words for which the next word followed later than 300 ms (n = 587) or earlier than 300 ms (n = 2908). The TRFs for these two groups of function words differed significantly, suggesting that the positive component at 200 ms is primarily due to short function words (see Figure 4-3). This finding thus suggests that the TRF to function words might be biased by a response to the subsequent word boundary.

### Across story

To confirm that responses to linguistic features are consistent across speaker and story, we verified whether linguistic speech representations have added value when the model was trained on DKZ and used to predict brain responses to other stories. Except for DWZ_1 (p=0.194) and DOL (p=0.083), a significant increase in prediction accuracy is seen when the linguistic speech representations are added (AEDV_1: p < .001, effect size=1.035; AEDV_2: p=0.013, effect size=0.466; DWZ_2: p=0.03, effect size=0.431; Eline: p < .001, effect size=0.799; Figure 5.A).

**Figure 5:**
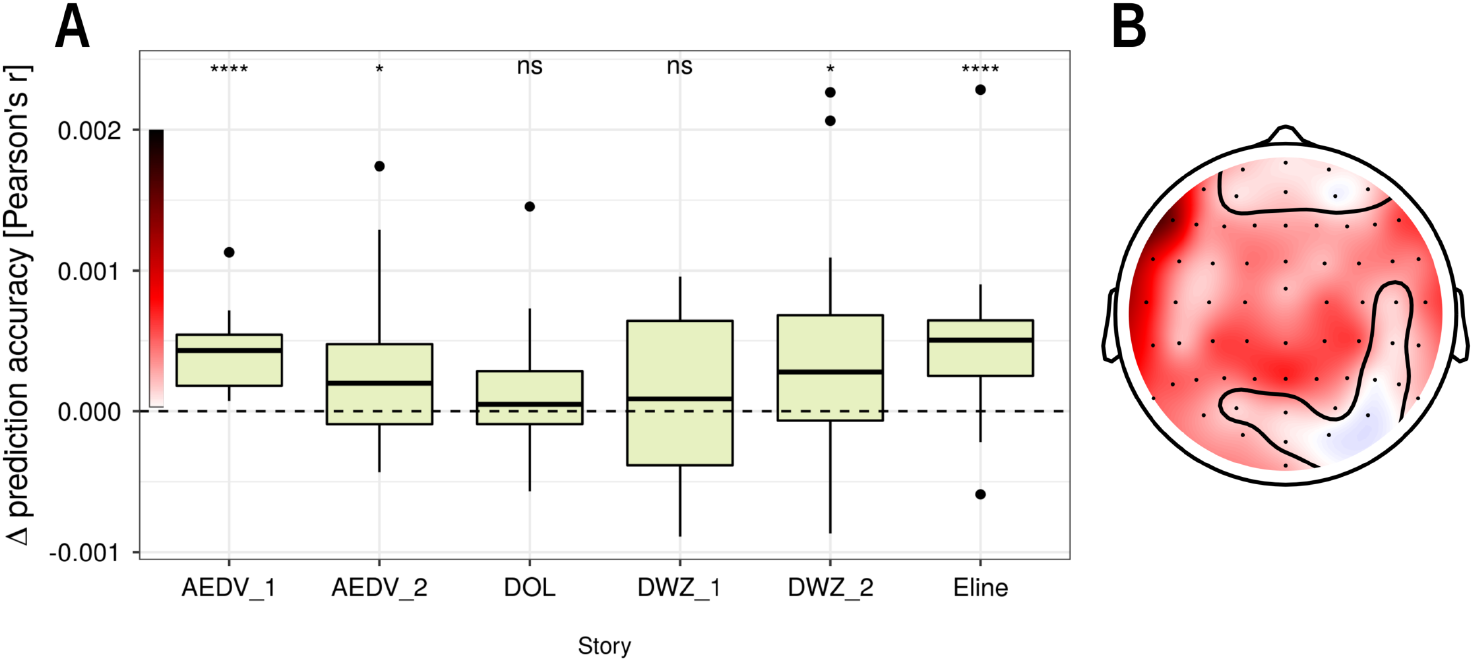
Added value of linguistic speech representations across story: Panel A: Increase in prediction accuracy (Pearson’s r) averaged across all sensors of the model including the linguistic representations compared to the model which only includes acoustic and speech segmentation properties of the speech. Panel B: The increase in prediction accuracy, averaged across stories, in sensor space. A cluster-based permutation test resulted in one large cluster encompassing almost all sensors; the channels which were *not* included in the cluster are encircled.

We observed that the added value of linguistic speech representations varied across the different stories. To determine whether or not this variation across the different stories was systematic, we identified the best LMM, using the predictors: story identity, presentation order and presentation distance (i.e., number of intervening stories between test story and DKZ). The latter two were included to capture fluctuations in subjects’ mental state over time. The Buildmer toolbox determined that the best LMM contains only the factor for story identity. A LMM using story identify (AIC=-1231.0) significantly improves the model fit compared to a model with only the random effect (AIC=-1229.6; Chi-square test comparing the two models returned a p-value of 0.04358). Based on the restricted maximum likelihood, the model fit did not improve when we included the presentation order (AIC=-1229.5), presentation distance ( AIC=-1230.2) or a combination of the latter two (AIC=-1228.8). We also verified this result with the Chi-square tests: the model fit did not improve when presentation order, distance, or a combination of the latter two were included on top of the story identify. This analysis shows that only the factor story identity improves the model fit when explaining the variance of the added value of linguistic speech representations among the different stories.

From the above analyses, we infer that there is some intrinsic variability among the stories as to how well the trained model generalizes. However, the current study does not have enough data points to systematically investigate which features of the story and/or speaker characteristics are the source of this variability. Such an investigation would require systematically varying those features, whereas our study only employed a fixed set of 6 different test stories. Curiously, the two stories with non-significant generalization were spoken by the same narrator as the training story, and there did not seem to be a distinguishing feature to identify those two stories and setting them apart from the others (see Table 3 and Table 1).

## Discussion

We evaluated which linguistic representations are tracked over and beyond acoustic and speech segmentation representations in EEG. Additionally, we showed that the tracking of linguistic representations is similar across stories.

### Reliable tracking of linguistic representations at the phoneme level

Both phoneme surprisal and cohort entropy had a significant added value compared to the acoustic and other linguistic representations, demonstrating that these effects, previously shown with magnetoencephalography (MEG) (Brodbeck et al., 2018), can be measured with EEG, and suggesting that both representations contribute independently to explaining neural responses.

Brodbeck et al. (2018) reported significantly different latencies for phoneme surprisal and cohort entropy respectively around 114 ms and 125 ms. We did not observe a corresponding difference in latency here. Additionally, Brodbeck et al. (2018) reported that the anatomical regions of the responses to these speech representations did not significantly differ. In our results, the topographic response of phoneme surprisal and cohort entropy in a time window around 100 ms did not significantly differ either, suggesting a similar neural source configuration (Table 3-2).

A difference with the neural responses in the study by Brodbeck et al. (2018) is that in our study, the neural responses show more than one prominent peak. While Brodbeck et al. (2018) did not elaborate on later activity in the TRFs, we observed a distinct negativity around 250 ms for central channels. This negativity did not significantly differ in latency or topography between the two linguistic representations. However, there was some indication that phoneme surprisal and cohort entropy might be associated with topographically different responses around 400 to 500 ms (Figure 3; Table 3-1) suggesting different underlying neural source configurations. This difference in topography is consistent with the interpretation that the two representations represent distinct speech processing stages. As Brodbeck et al. (2018) suggested, phoneme surprisal might reflect a measure of phoneme prediction error which is used to update the active cohort of lexical items, and cohort entropy might reflect a representation of this cohort of activated lexical items. The difference between the two studies might be due to the difference in modality (EEG vs. MEG) or task (for example, in the MEG study some trials were repeated whereas here all stimuli were presented only once).

### Reliable tracking of linguistic representations at the word level

At the word level, only word surprisal and word frequency had a significant added value compared to each other and acoustic and speech segmentation representations.

Word surprisal was identified as a significant predictor which is in line with the previous literature (Weissbart et al., 2020; Koskinen et al., 2020). Although word frequency and word surprisal are correlated, there is an added value of word frequency over and beyond word surprisal (and vice versa). The neural responses to both linguistic representations show a negativity around 400 ms in central parietal areas, analogous to the typical N400 response derived in ERP studies. Interestingly, we observed a significant difference in topography between word surprisal and frequency (Figure 3; Figure 3-4) suggesting that the responses are due to different neural source configurations. It is hypothesized that the N400 response reflects multiple processes, including activation of lexical items in memory, and the semantic integration of the word into its context (for a review Lau et al. (2008) and Kutas and Federmeier (2011)). Our findings suggest that word surprisal and word frequency might index slightly different processes reflected in the N400 during language comprehension: the response to word frequency might be related primarily to the activation of lexical items, as a word with a higher frequency is easier to access in long term memory, while word surprisal might reflect a combination of lexical activation and semantic integration.

Although previous studies reported an added-value of word entropy and word precision (Willems et al., 2016; Weissbart et al., 2020), those predictors did not significantly improve the prediction accuracy in our data. Using functional magnetic resonance imaging (fMRI), Willems et al. (2016) reported significant responses to word entropy. However, word entropy was modeled as the uncertainly of the next word while in our study it was defined as the uncertainty of the current word. If the effect observed in fMRI reflects brain activity related to predicting the next word, as suggested by Willems et al. (2016), then we might not expect an effect of current-word entropy in EEG, as the corresponding brain activity might have occurred on the previous word. Another important difference is the imaging modality; possibly, the more distributed parietal and frontal sources associated with entropy are less visible in EEG. Moreover, our EEG methodology assumes strictly time-locked effects. Thus, if an effect is not strictly time-locked, it might be detected in fMRI but not in EEG.

We also did not observe a significant effect of word precision, in contrast to Weissbart et al. (2020). These divergent results might be explained by a difference in methodology: We focus on a significant added value in prediction accuracy, while Weissbart et al. (2020) determined the significance of the TRF. Because different speech representations are derived from the same speech signal, they are usually correlated. Therefore, a speech representation that does not significantly improve predictions can nevertheless obtain a significant TRF due to its correlation with other significant speech representation. By testing prediction accuracies, we evaluated the different speech representations more conservatively. It is, of course, possible that word precision is associated with a real effect but only provides very little non-redundant information, and such a small effect might not have been detected in our study.

### No significant neural tracking of linguistic representations at the contextual level

Semantic dissimilarity did not show a significant added value over and beyond acoustic and speech segmentation features. This is in contrast to a result by Broderick et al. (2018) showing that when listening to narrative speech, semantic dissimilarity is tracked by the brain. To address this discrepancy, a more detailed analysis was performed, which showed that semantic dissimilarity does provide added value over and beyond content words, but fails to do so when also controlling for acoustic and speech segmentation features. Similarly, Dijkstra et al. (2020) reported that no added value of semantic dissimilarity was seen after controlling for the acoustic envelope and content word onsets. On the other hand, brain responses seem to be sensitive to semantic dissimilarity under some conditions, as a study of sentence reading found N400-like effects of semantic dissimilarity (Frank and Willems, 2017), and a parallel fMRI investigation, where the participant listened to fragments of audiobooks, localized activity correlated with semantic dissimilarity in non-auditory brain areas (Frank and Willems, 2017). One possibility that might account for all these observations is that semantic dissimilarity might be highly correlated with acoustic properties of speech, and thus would not survive correction for acoustic predictors.

### Neural tracking of linguistic representation is independent of the word class

The differentiation between content and function words for word surprisal and word frequency did not have an added value. Similar to the findings of Frank et al. (2015) and Brennan and Hale (2019), this suggests that the neural response to these linguistic representations depends on the variation between subsequent words, independent of their word class.

### Neural tracking across stories

When the model trained on one story is applied to another, an added value of the linguistic representations is seen in 4 out of 6 stories. This is not just due to random variability, as the observed prediction accuracies differ significantly across stories, and presentation order does not explain this variability. A possible explanation is that some stories may be more appealing than others due to the story content, influencing the signal to noise ratio (SNR) of the EEG responses. However, our dataset was too limited to investigate the effects of speaker and story characteristics systematically. Future studies might look into this matter by collecting data with a more varied sample of different stories to generate the required variability in speaker and story characteristics.

### Caveats

We want to emphasize two points of concern when interpreting our results. Firstly, the *unique* contribution of linguistic representations to the EEG signals is small, when compared to the acoustic representations. This is likely due to a combination of factors. Speech is a spectro-temporally complex and broadband signal capable of eliciting activity in large parts of the auditory cortex (Hullett et al., 2016). In contrast, the linguistic representations model much more specific processes, likely eliciting activity in correspondingly smaller regions of cortex, leading to less current that can still be measured at the scalp EEG. Moreover, our tests are inherently conservative: linguistic representations by themselves might be able to explain a much larger amount of the variance in the EEG signal, but our tests quantify the variability *uniquely* attributable to linguistic representations, not counting variance that is shared with acoustic (and other linguistic) representations. While the absolute magnitude of the responses to linguistic representations is thus expectedly small, we found that they can be reliably detected across subjects, with moderate to large effect sizes.

Secondly, we did not compare an intelligible to an unintelligible condition. Consequently, our results do not directly provide evidence that these representations are related to speech intelligibility. For some representations, such a relationship has been shown by previous investigations, e.g. TRFs to semantic dissimilarity (Broderick et al., 2018) and the phoneme level variables (Brodbeck et al., 2018) flattened when the speech was not understood. For other representations, such a test is still outstanding. As discussed in the introduction, a correlation with intelligibility is not by itself evidence that a representation can measure intelligibility. However, we would argue that the combination of both – a correlation with intelligibility *and* predicting brain signals beyond acoustic properties – can make a stronger case that the neural tracking of linguistic speech representations is an index of language processing rather than a side-effect of acoustic speech representations.

## Conclusion

Linguistic representations explain the neural responses over and beyond acoustic responses to speech. We found significant neural tracking of phoneme surprisal, cohort entropy, word surprisal and word frequency over and beyond the tracking of the speech’s acoustic properties and speech segmentation. This was not observed for word entropy, word precision and semantic dissimilarity. In this paper, we showed the importance of controlling for acoustic and speech segmentation properties of the speech when estimating the added value of linguistic representations.

Additionally, we were able to predict neural responses to speakers and stories unseen during training. This suggests that the processing of these linguistic representations is independent of the presented content and speaker and, therefore, show evidence that higher stages of languages processing are modelled. Therefore these linguistic representations show promise for a behaviour-free evaluation of the speech intelligibility in audiological and other clinical settings.

## Acknowledgements

We would like to thank Bernd Accou for his continuous effort to collect data and for creating the alignment of phonemes and words to the speech signal. Furthermore, we would like to thank Hugo Van hamme for allowing us to use the constructed language models. Additionally, we want to thank all members of the ExpORL ISIFIT team for their weekly guidance.

## Extended Data

### Extended data regarding Figure 2

**Figure 2-1:**
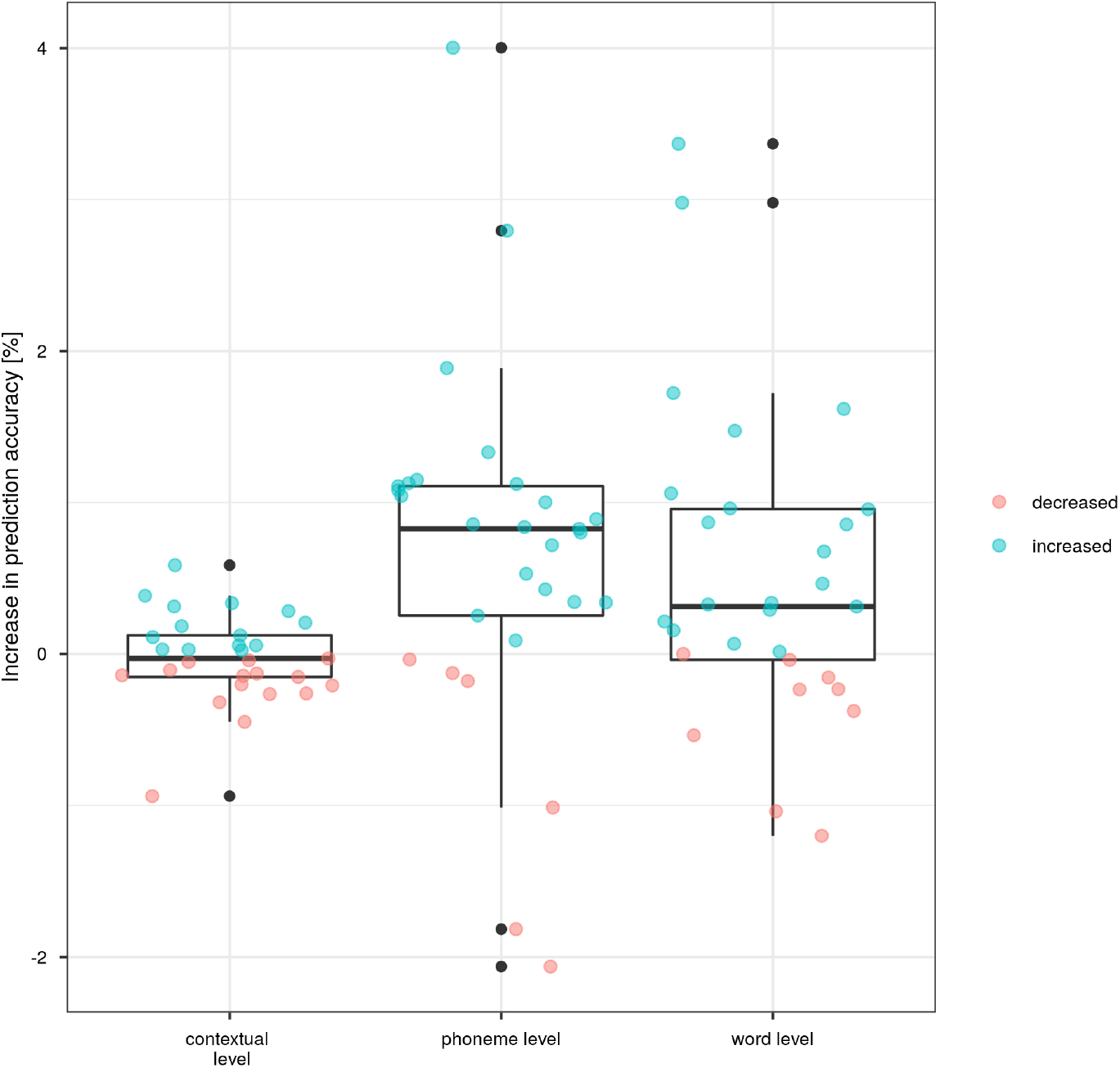
Increase in prediction accuracy when the linguistic representations are added to a model that includes acoustic and speech segmentation properties at a contextual, phoneme, and word level. If the prediction accuracy increases, the data point is shown in blue; otherwise, the line is red.

### Extended data regarding Figure 3

**Table 3-1:**
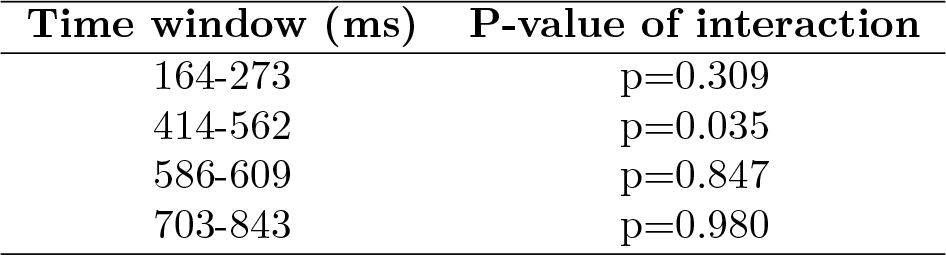
Intersections of the significant time windows of phoneme surprisal and cohort entropy with the p-value of the interaction term between sensor and speech representation obtained via the method proposed by McCarthy and Wood (1985) for a central channel selection.

**Figure 3-1:**
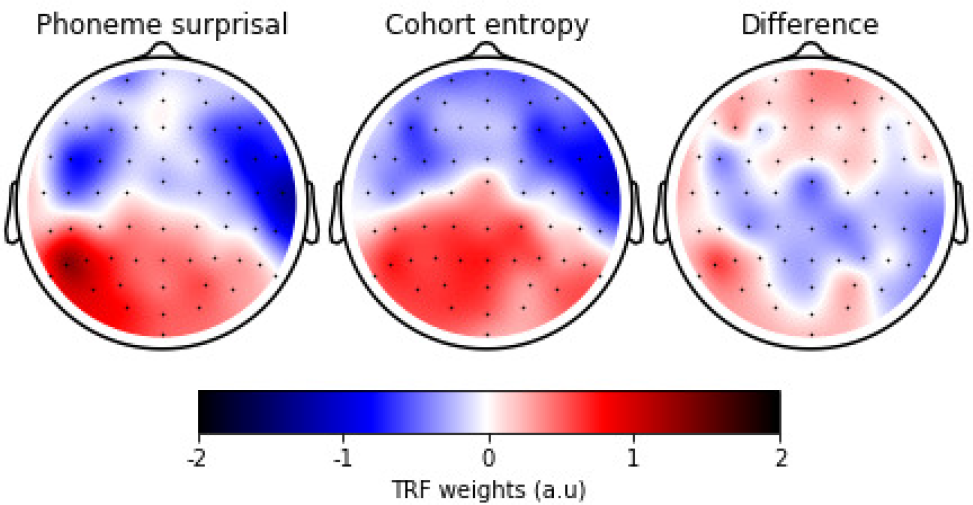
The normalized TRF-weights averaged across participants within the time window 414 ms to 562 ms for phoneme surprisal (left) and cohort entropy (middle) and the resulting difference (left).

**Table 3-2:** Intersections of the significant time windows of phoneme surprisal and cohort entropy with the p-value of the interaction term between sensor and speech representation obtained via the method proposed by McCarthy and Wood (1985) for a frontal channel selection.

| Time window (ms) | P-value of interaction |
| --- | --- |
| 102-164 | p=0.263 |
| 367-461 | p=0.634 |
| 773-914 | p=0.994 |

**Table 3-3:**
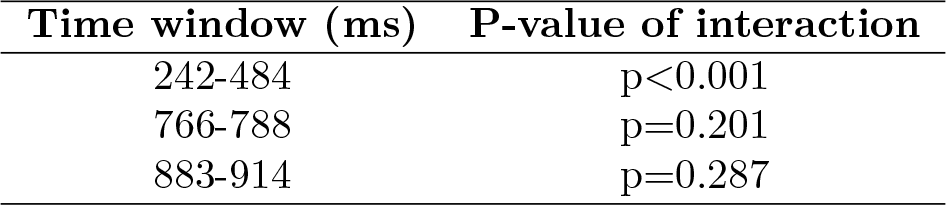
Intersections of the significant time windows of word surprisal and word frequency with the p-value of the interaction term between sensor and speech representation obtained via the method proposed by McCarthy and Wood (1985) for a central channel selection.

**Figure 3-2:**
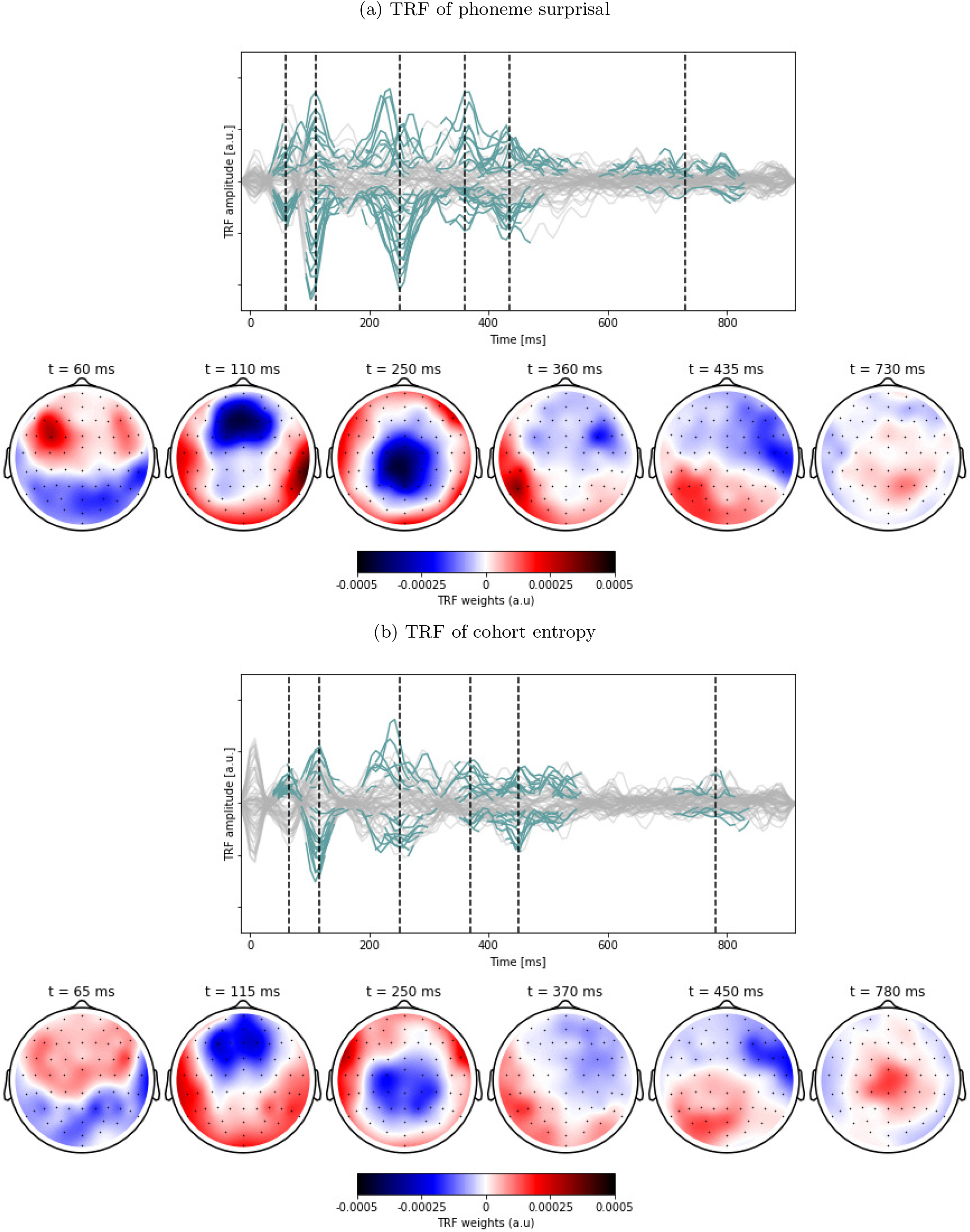
The average TRF to the linguistic predictors at the phoneme level. The channel responses over time which are significantly different from zero, are annotated in blue. The insets below show the topographic responses at the peak latencies annotated with the dashed vertical lines.

**Figure 3-3:**
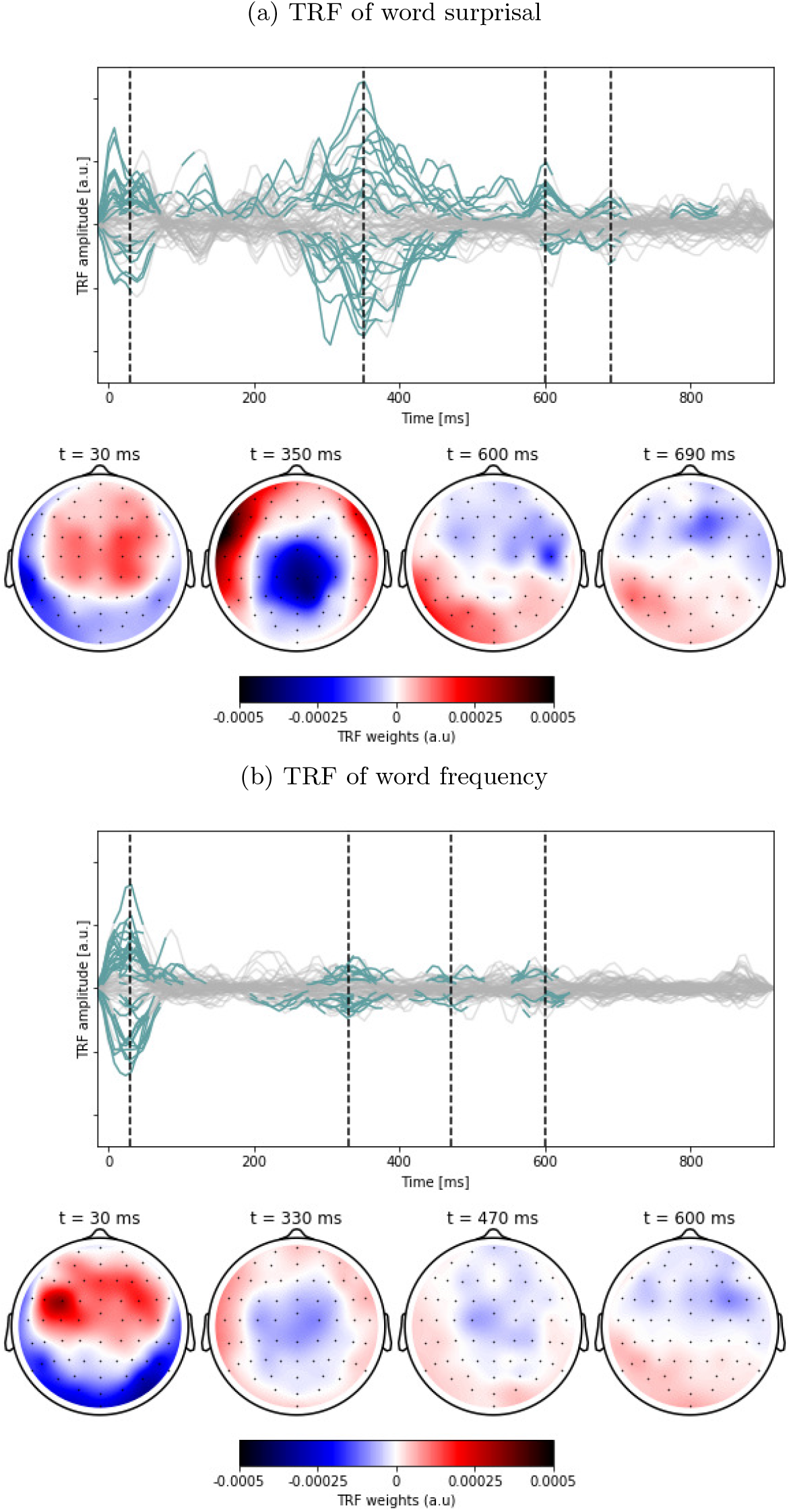
The average TRF to the linguistic predictors at the word level. The channel responses over time which are significantly different from zero, are annotated in blue. The insets below show the topographic responses at the peak latencies annotated with the dashed vertical lines.

**Table 3-4:**
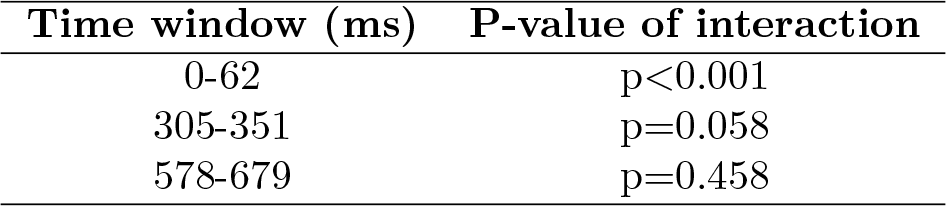
Intersections of the significant time windows of word surprisal and word frequency with the p-value of the interaction term between sensor and speech representation obtained via the method proposed by McCarthy and Wood (1985) for a frontal channel selection.

**Figure 3-4:**
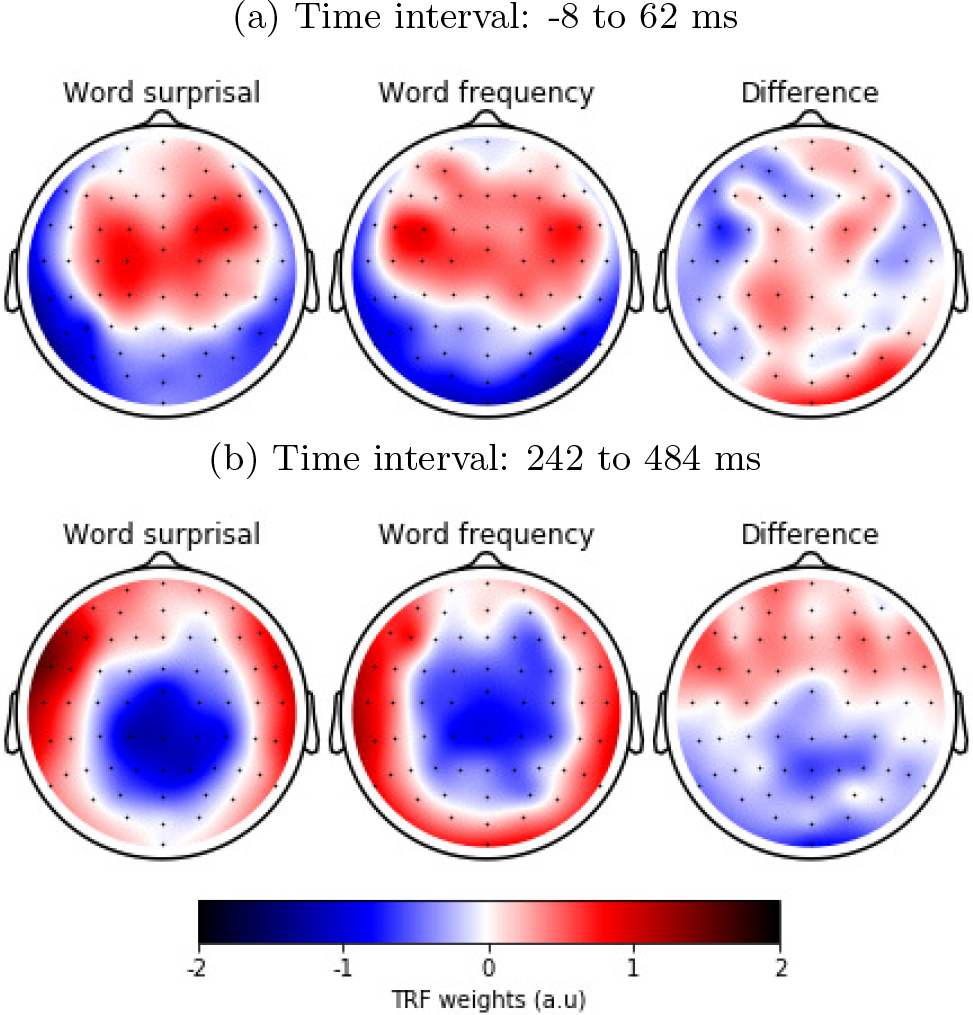
The normalized TRF-weights averaged across participants within 2 different time windows for word surprisal (left) and word frequency (middle) and the resulting difference (left).

**Figure 3-5:**
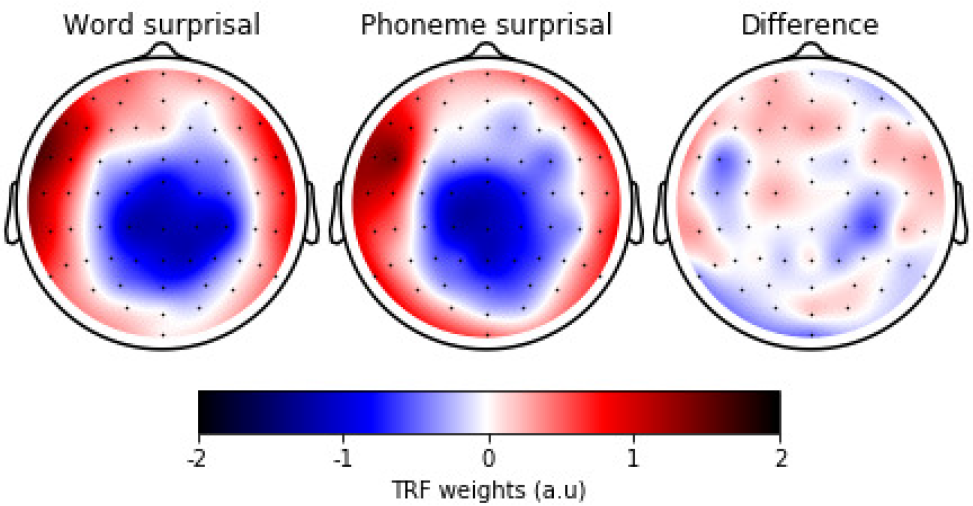
The normalized TRF-weights averaged across participants within the time window 242 ms to 531 ms for word surprisal (left), 164 ms to 343 ms for phoneme surprisal (middle) and the resulting difference (left).

### Extended data regarding Figure 4

**Table 4-1:** Time windows of prominent peaks in the response to content and function words with the p-value of the interaction term between sensor and speech representation obtained via the method proposed by McCarthy and Wood (1985).

| Time window (ms) | P-value of interaction |
| --- | --- |
| -16-86 | $p < 0.001$ |
| 148-164 | $p < 0.001$ |
| 227-25 | $p < 0.001$ |
| 297-375 | $p < 0.001$ |
| 523-547 | $p = 0.436$ |

**Figure 4-1:**
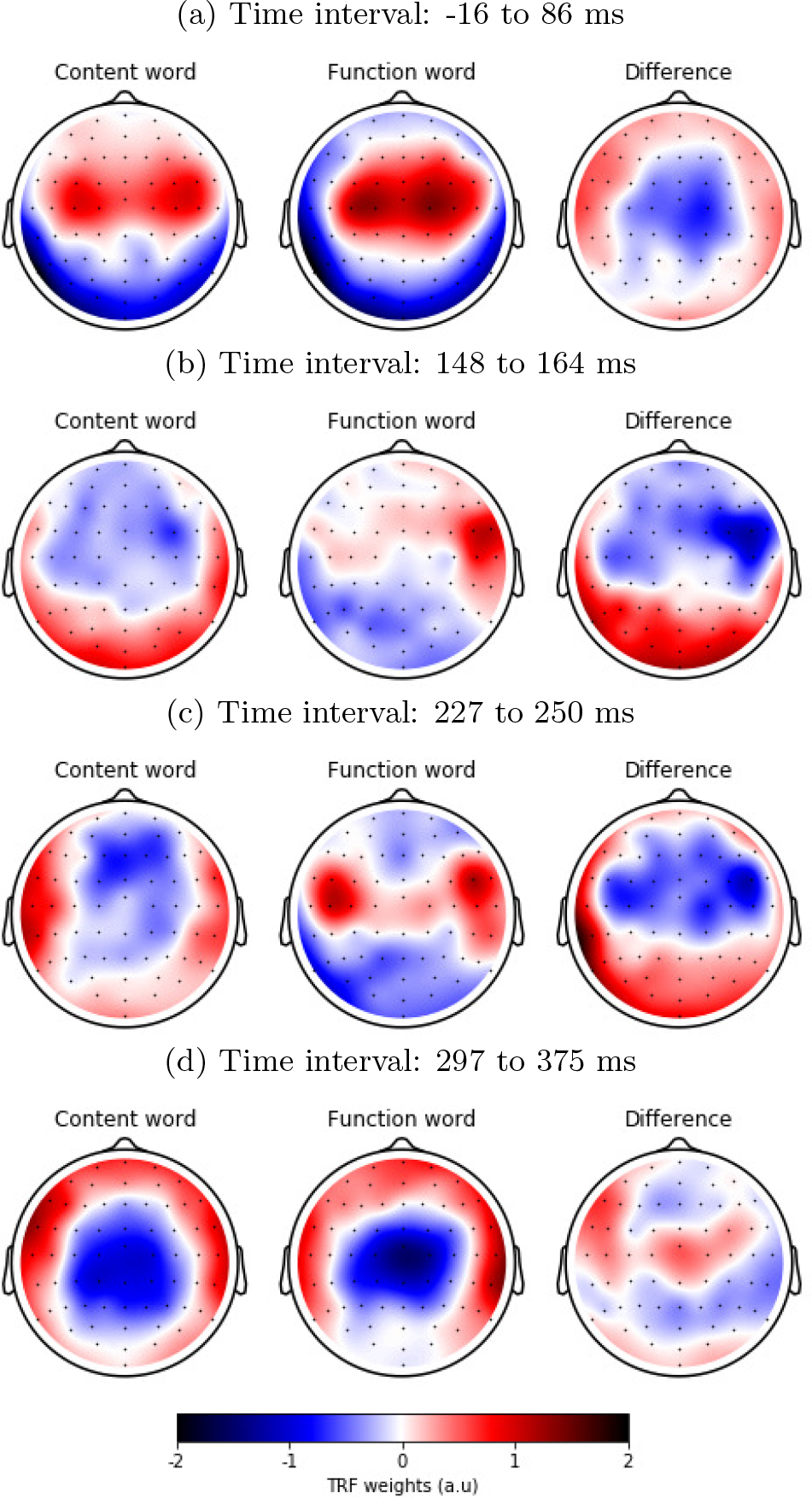
The normalized TRF-weights averaged across participants within 4 different time windows for the response to a content word (left) and a function word (middle) and its resulting difference (left).

**Figure 4-2:**
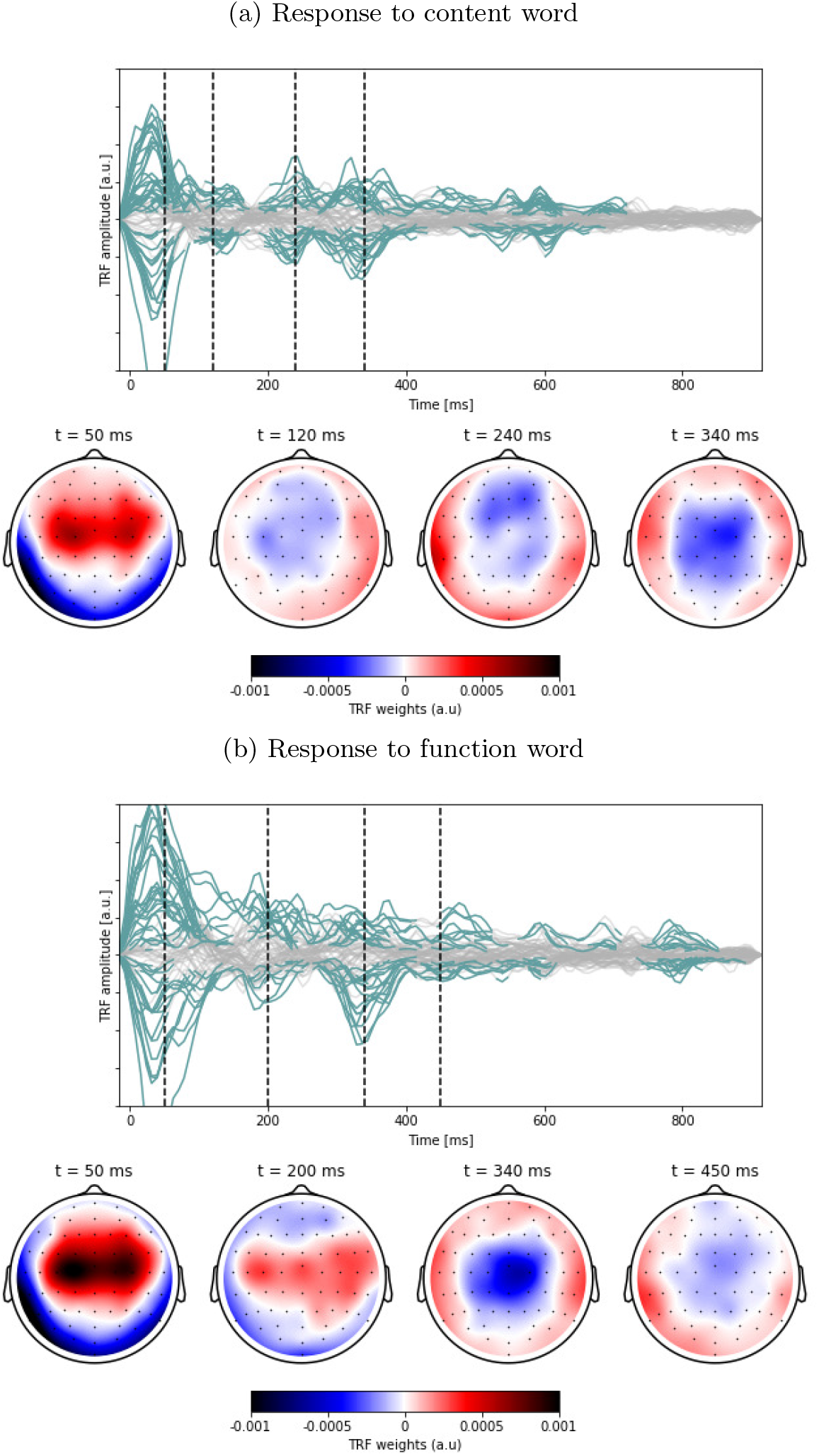
The average neural response to a content (top) and function word (below). The channel responses over time which are significantly different from zero, are annotated in blue. The insets below show the topographic responses at the peak latencies annotated with the dashed vertical lines.

**Figure 4-3:**
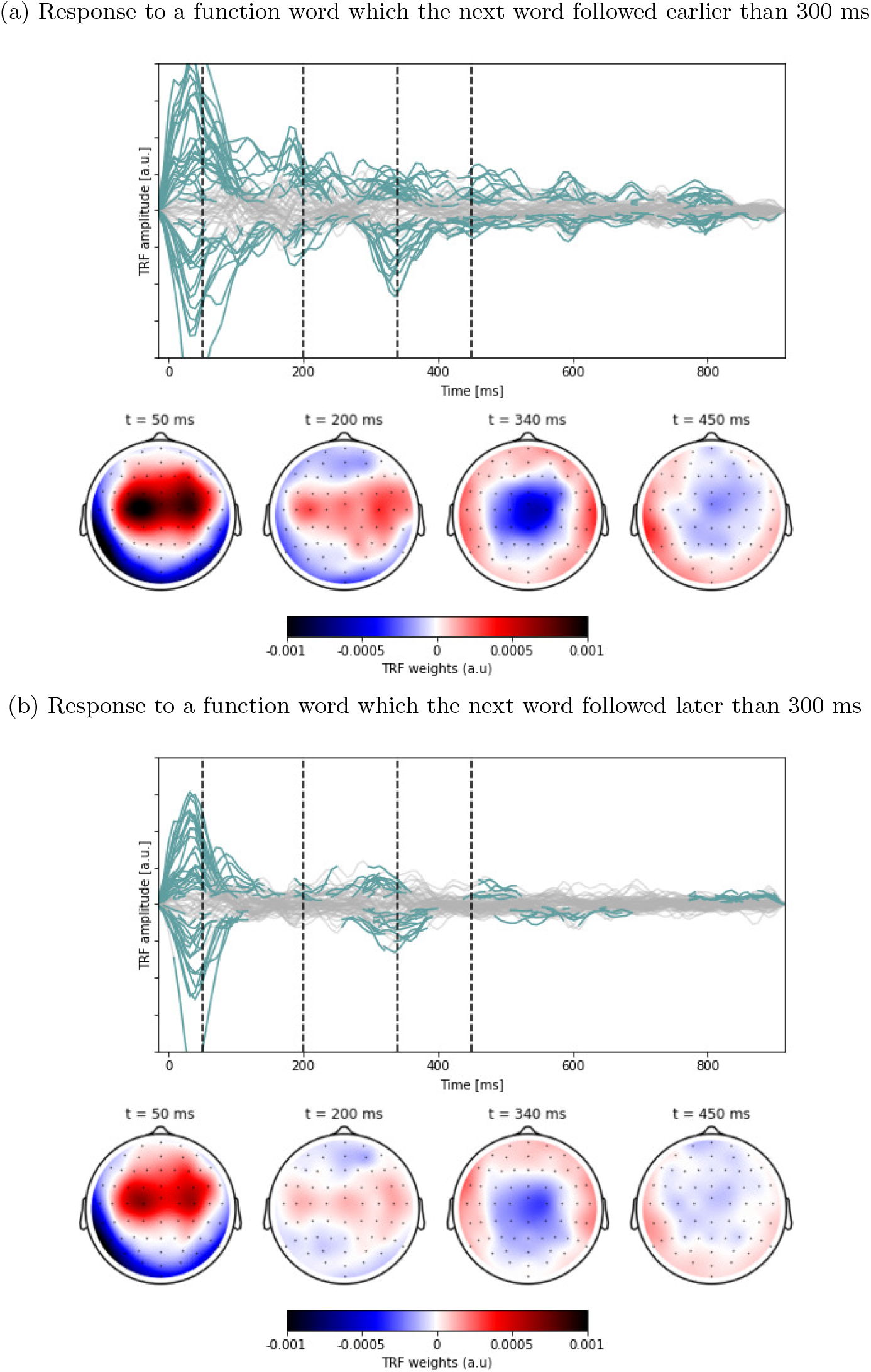
The average neural response to a function word which the next word followed earlier than 300 ms (top) and later than 300 ms (below). The channel responses over time which are significantly different from zero, are annotated in blue. The insets below show the topographic responses at the peak latencies annotated with the dashed vertical lines.

**Figure 4-4:**
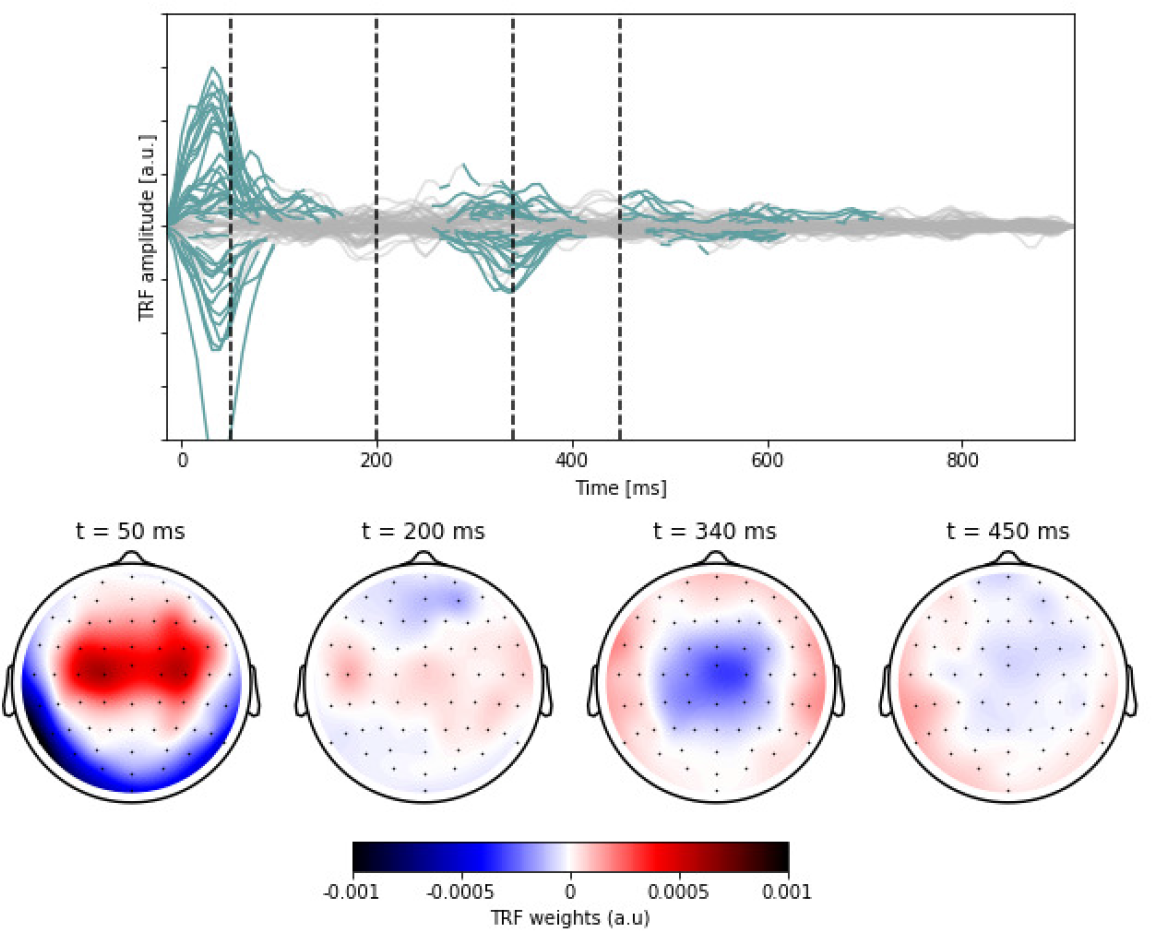
The average neural response to word onsets. The channel responses over time which are significantly different from zero, are annotated in blue. The insets below show the topographic responses at the peak latencies annotated with the dashed vertical lines.

### Extended data regarding Figure 5

**Figure 5-1:**
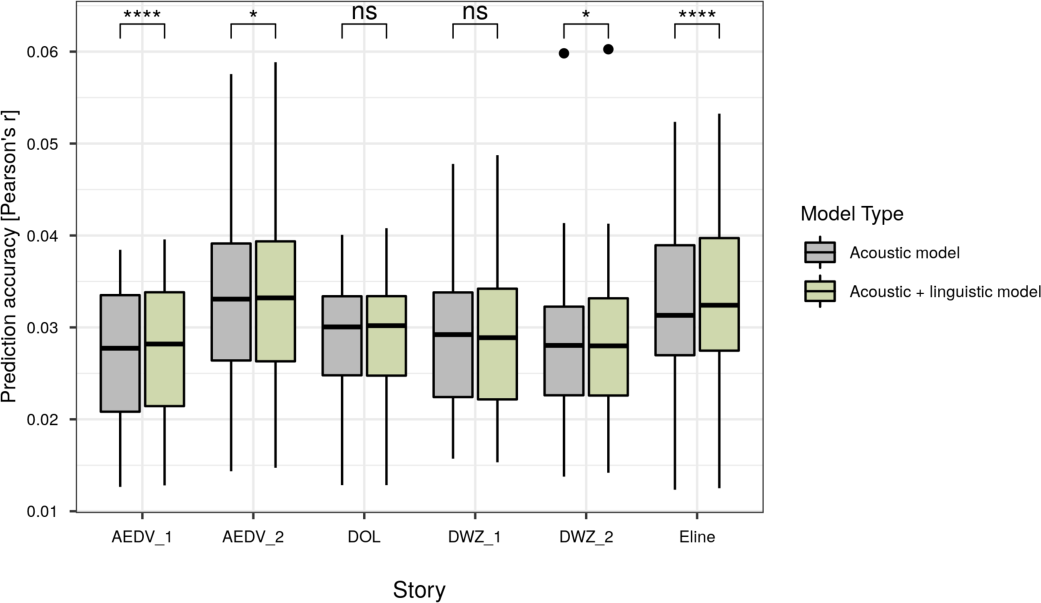
Comparison of the prediction accuracies of the acoustic model (grey) and the acoustic model with linguistic predictors (green) for each story.

